# The Panama Canal after a century of human impacts

**DOI:** 10.1101/777938

**Authors:** Jorge Salgado, María I. Vélez, Catalina González-Arango, Neil L. Rose, Handong Yang, Carme Huguet, Juan Camacho, Aaron O’Dea

**Author notes:** corresponding author: Jorge Salgado.

## Abstract

Large tropical river dam projects are set to accelerate over the forthcoming decades to satisfy growing demand for energy, irrigation and flood control. When tropical rivers are dammed, the immediate impacts are well studied, but the long-term (decades-centuries) consequences of impoundment remain poorly known. Here, we gather historical and paleoecological data from Gatun Lake, formed by the building of the Gatun Dam (Panama Canal, Panamá) over 100 years ago, to reconstruct the limnological evolution of the system in response to individual and linked stressors (river damming, forest flooding, deforestation, invasive species, pollution and hydro-climate). We found that after a century of dam construction parallels associated with the natural hydrological functioning of river floodplains persist. Hence, hydrology remains the most important temporal structural factor positively stimulating primary productivity, deposition of new minerals, and reduction of water transparency during wet periods. During dry periods, clear water and aerobic conditions prevail and nutrients transform into available forms in the detrital-rich reductive sediments. We highlight the importance of climate change as an ultimate rather than proximate anthropogenic factor for sustainable management options of tropical dams.

## Introduction

The tropics contain more than 4 million bodies of freshwater^1^ and many large tropical rivers have been dammed for water management, commerce and energy production^2^. Projections show a three-fold increase over the forthcoming decades in the number of large dammed (>15 m high) tropical river projects^3–5^. When rivers are dammed, the immediate impacts are well studied^3^. For instance, dams transform carbon cycling^6, 7^, alter river networks by creating artificial reservoirs^3, 8^, alter natural patterns of sediment transport^8^, impede upstream-downstream movement of fish^3^ and modify water quality and productivity^3, 8^. Yet almost nothing is known about the long-term impacts and limnological evolution of impoundment of highly diverse, tropical rivers. Rivers in the neotropics (the tropical areas of North, Central and South America) are, in particular, poorly studied^8, 9^ with almost no long-term continuous time-series of data for multiple variables^10^. This constrains our understanding on how these aquatic systems can be sustainably managed in the long-term. In this study we combined exemplary historical and paleoecological approaches to provide a continuous long-term record of lake hydrology and geochemistry and the responses of different aquatic biological groups associated with the iconic Panama Canal.

In 1913 the Chagres River was dammed, forming Gatun Lake on which the Panama Canal is located (Fig. 1). At the time, Gatun Lake was the first neotropical large dam and was the largest man-made lake in the world with a surface area of 425 km^2^. In contrast, most neotropical dams are no older than a quarter of century^8, 10^ making Gatun Lake a unique opportunity to reveal long term dynamics in a tropical dam system. Moreover, Gatun Lake has extensive environmental and ecological records thanks to monitoring and research programs established by the Panama Canal Authority (ACP for its acronym in Spanish) and the longstanding presence of the Smithsonian Tropical Research Institute (STRI) in Panama. Combining these records that are unavailable for other old dams, with paleoecological data offer therefore an unrivalled opportunity to explore the impacts of damming on a tropical river over a period of more than 100 years.

**Figure 1.**
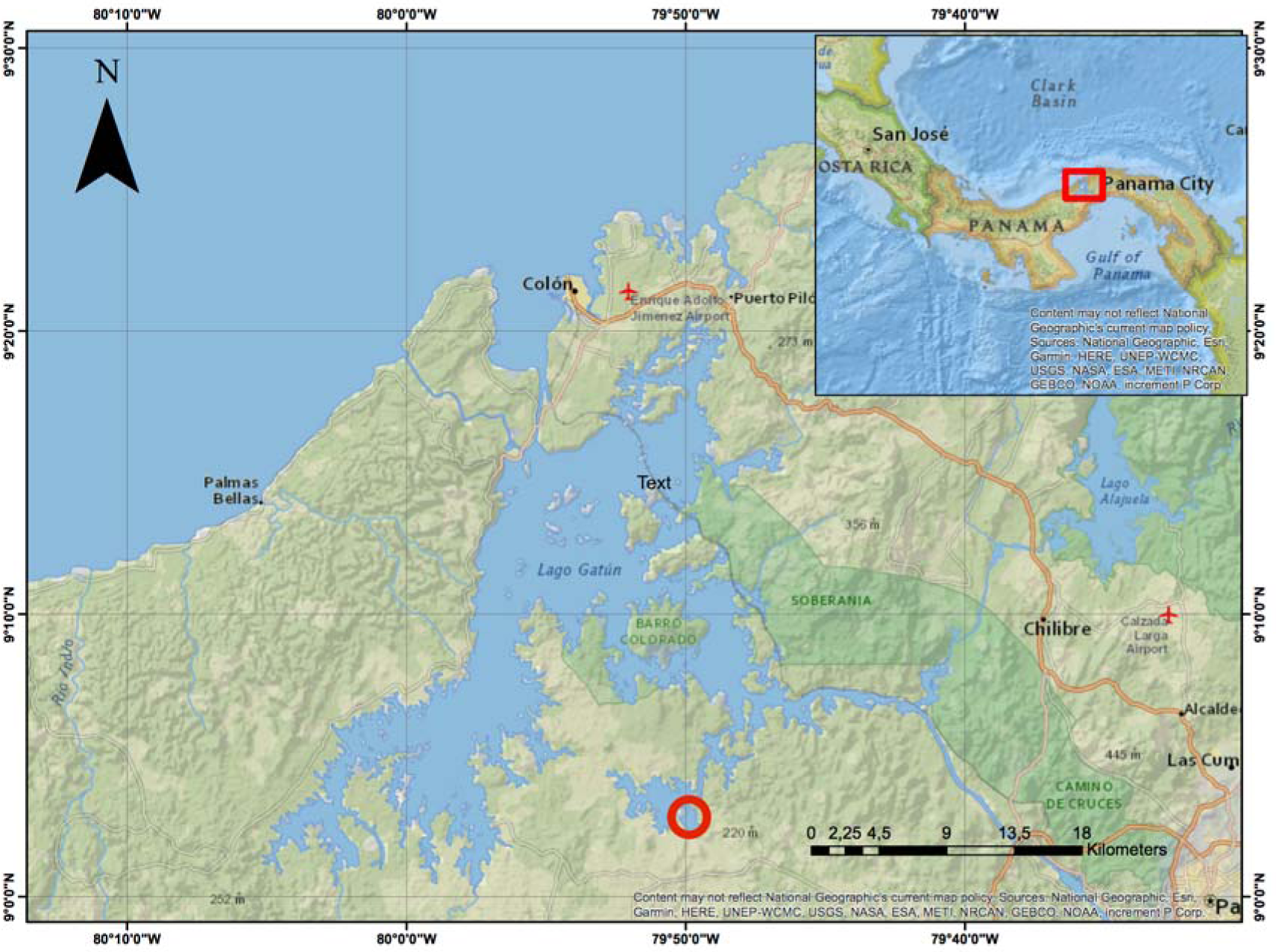
Map of the River Chagres watershed. The artificial Gatun and Alajuela lakes and the connecting River Chagres are indicated in blue. A red circle shows the coring location of LGAT1 core. Natural protected areas are shown in dark-green.

In this study, we present sedimentary data of Gatun Lake to build a biological and environmental chronological sequence of changes and combined it with historical water quality records (secchi depths, pH, conductivity and dissolved oxygen) available from 1969 to 2013, and on river annual flow and climate (precipitation) records available from 1930 to 2013. The aim was to: (1) reconstruct the limnological evolution of the Chagres River landscape (including pre-impoundment times); and (2) assess individual and linked effects of river damming, forest clearance and flooding, the introduction of invasive species, water quality variation and climate on this aquatic landscape.

## Results

### Environmental history of Gatun Lake

The canal region experiences a seasonal tropical monsoonal climate. Mean annual temperature across the canal area is 26 °C and mean monthly temperature varies by just 1°C annually^11^. Rainfall ranges from around 1750 mm year^−1^ mean annual precipitation on the Pacific coast, to 4000 mm year^1^ on the Caribbean coast^12^. Records beginning in the late-1920s on the Smithsonian’s Barro Colorado Island (BCI) in the canal show that annual rainfall has varied considerably between years, apparently related to ENSO conditions^13^.

Gatun Dam led to flooding of large areas of tropical forest^14^, an increase in sediment accumulation^15^ and, watershed deforestation^16, 17^ for agriculture, in addition to the operation of the canal and urban expansion^17^. Almost 20 years later, another portion of rainforest was flooded across the river headwaters by the construction of a second dam, Lake Alajuela, built to regulate the stochastic hydrology of the Chagres River^15^ (Fig. 1). The consequences of forest flooding have yet to be quantitatively assessed, but is likely that the expansive areas of flooded allochthonous organic material would have contributed considerable quantities of DOC into the newly formed lake^6, 7^ facilitating large emissions of greenhouse gases to the atmosphere, even decades after construction of the dam, as has been shown in other tropical dams^7, 18^.

Salinity of Gatun Lake has been historically below 0.2 ppt^19^. However, increases in salinity above the United States Environmental Protection Agency (EPA) drinking water standards (>3.0 ppt) have been reported in the Miraflores locks during the dry season since the completion of the Canal^20^. Furthermore, recent (2004-2015) expansion of the canal has resulted in higher water demands to operate a new set of larger locks resulting in more salt water penetrating the lake, especially during the dry season^20^. The consequences of this emergent salinization of Gatun Lake continue to be debated but observations of brackish water fauna, such as the non-native North American Harris mud crab (*Rhithropanopeus harrisii*)^21^ and the Iraqi crab (*Elamenopsis kempi*)^22^, and increases in observations of marine fish have already been reported in the lake^23^.

Gatun Lake has also been subjected to introductions of several nuisance aquatic species. The Asian macrophyte *Hydrilla verticillata* was first recorded in the lake around the 1930s, which rapidly dominated the lake after it was filled^24^. Introduction of the peacock bass (*Cichla ocellaris*) to the lake in the late-1960s caused a major ecological reorganization associated with dramatic declines in native littoral planktivorous fish species^25^, and these impacts endure 45 years later^23^. Other known introductions have included the Asian bryozoan *Asajirella gelatinosa*^26^, the Asian *Melanoides* (Thiara) *tuberculata* and, the South American apple (*Pomacea bridgesii*) snail^27^. There are likely myriad other introductions that have been undocumented or that are projected to occur under the new canal expansion^28^.

### Paleoecological data

We focused our paleoecological analysis on three biological groups from which fossil remains are commonly well preserved in lake sediments: macrophytes, invertebrates (bryozoans, chironomids, cladocerans and mollusks) and diatoms. As a measure of temporal environmental change we combined information on the variation of single trace elements (potassium [K]–river erosion; calcium [Ca]–salinity; lead [Pb], copper [Cu] and zinc [Zn]– pollution)^29^ and elemental ratios (iron/magnesium [Fe/Mn]–reductive conditions; titanium/calcium [Ti/Ca–detrital inputs])^29^. The carbon contribution of terrestrial plants, macrophytes, and bacteria-algae was assessed through *n*-alkanes C_15_–C_31_ biomarkers^30^. The *n*-alkanes information was also used to calculate the terrigenous aquatic ratio (TAR) and the submerged/floating aquatic macrophyte inputs vs. emergent/ terrestrial plant input ratio (Pmar-aq)^31^. A methane index (MI) ^32^ was obtained via archaea-glycerol dialkyl glycerol tetraethers [GDGTs]). Major zones of temporal change were assessed through clustering analysis and the links between biological and environmental variation were tested via multiple factor analysis–MFA^33^.

### Core chronology and sedimentation rates

A sediment core (87 cm long; LGAT1) was retrieved from a littoral area (9°2’49.58″N, 79°50’6.33″W) at a water depth of approximately 1 m (Fig. 1). Sediments were dated using the constant rate of ^210^Pb supply (CRS) model^34^ resulting in a chronology spanning the last c.150 years (Fig. 2). The pre-canal riverine conditions were contained within the 87-50 cm of the core and the post-canal lake conditions within the top 50 cm. The age model showed that sedimentation rates at post-canal times remained relatively uniform (each cm representing c. 8-15 years) until the early-2000s when rates almost doubled (Fig. 2).

**Figure 2.**
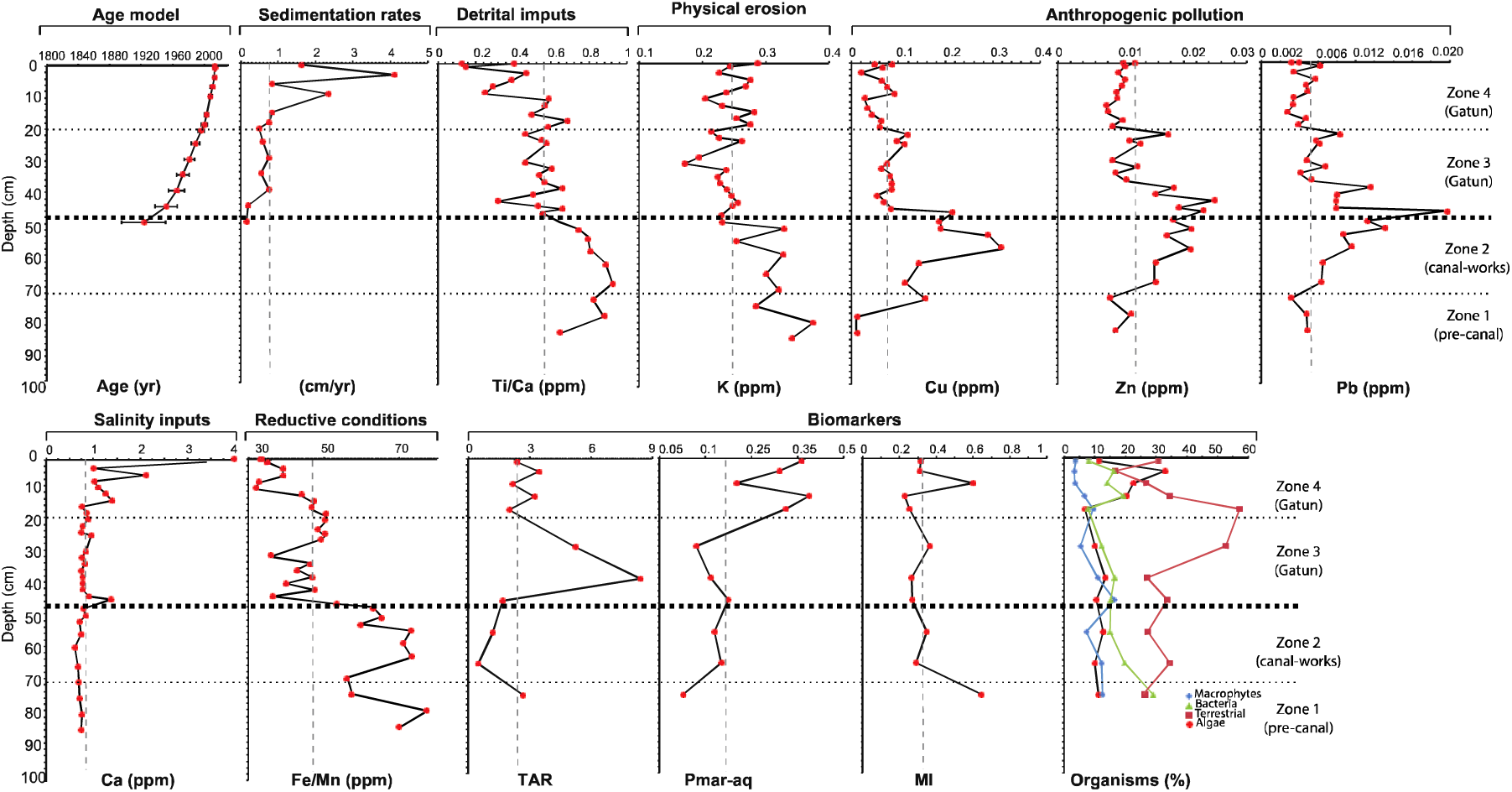
Sedimentary profile of the ^210^Pb age model (dates and ages and standard deviations are presented), sedimentation rates, selected geochemical elements and ratios, and biomarkers indices in LGAT1 sedimentary core. Terrigenous aquatic ratio–TAR; Methane Index –MI; the submerged/floating aquatic macrophyte inputs vs. emergent/ terrestrial plant input ratio–Pmar-aq. Major temporal zones of change determined by clustering analysis, corresponding to Zone 1 c. pre-1870, Zone 2 c.1870-1914, Zone 3 1923-1990, and Zone 4 1991-2013. A vertical grey dotted line indicates the mean value of each parameter.

### Long-term shifts in paleoecological proxies

Temporal variation on the selected geochemical and biomarker data is presented in Fig. 2. Sixteen macrophytes, 19 invertebrates and 81 diatom taxa (Figs. S1-S3) were identified throughout the sediment core and summarized into functional groups according to growth type (macrophytes), habitat preference (diatoms) and feeding mode (invertebrates) as follows (Fig. 3): *macrophytes* submerged; anchored-floating; free-floating and emergent; *invertebrates* filter-feeders (bryozoans), macrophyte/detrital (chironomids), shredders (Trichoptera cadis case larvae), benthic (chironomids), and grazers (mollusks and cladocerans); *diatoms* planktonic, benthic, littoral, aerophilous, and salinity-tolerant^35, 36^. Diatom benthic species were further sub-grouped into acidic, oligotrophic, benthic-mobile and eutrophic species^37–39^.

**Figure 3.**
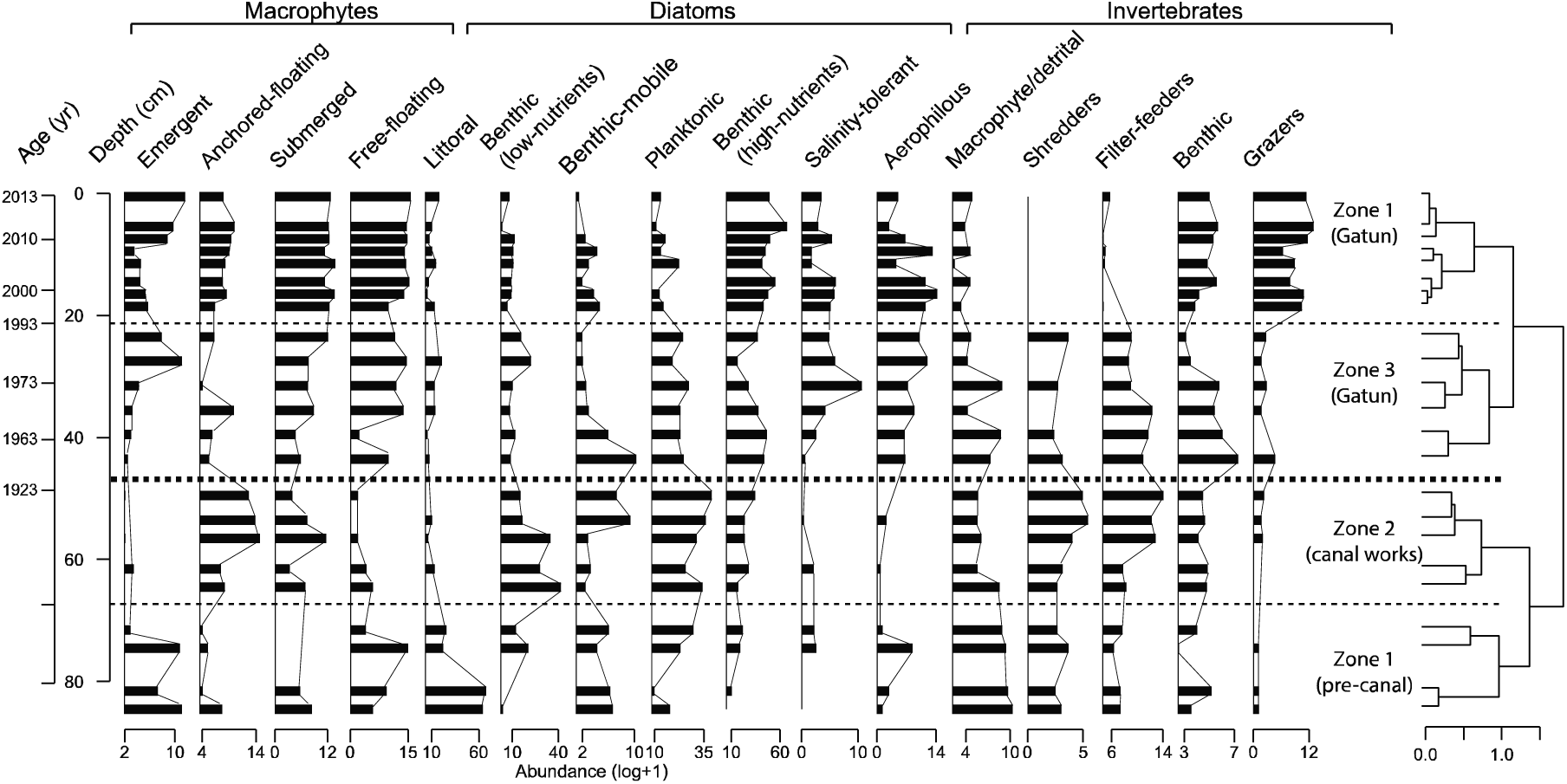
Sedimentary profile of the study biological functional groups in LGAT1 sedimentary core. Major temporal zones of change determined by clustering analysis are shown by dotted lines, corresponding to Zone1 *c.* pre-1870, Zone 2 *c.*1870-1914, Zone 3 1923-1990, and Zone 4 1991-2013.

Results uncover a dynamic aquatic history from before the formation of Gatun Dam by the Panama Canal to the present day. As expected, results describe over four main temporal zones of biological and environmental change a gradual transition from a river-governed system to a lake system (Figs. 2,3). During pre-canal times (Zone 1 c. pre-1870), the geochemical and biomarker data reflected swamp-like alluvial conditions characterized by reductive sediments (high MI index and Fe/Mn ratio respectively), low nutrient and acidic waters, and high detrital (Ti/Ca) inputs (Figs. 2,4). Before the creation of the Panama Canal, the Chagres River meandered through an alluvial floodplain of vast areas of swampy conditions^13, 40^. In agreement with those records, we found a prevalence of rushes and sedges, free-floating plant species (e.g. *Ludwigia sedoides, L. helminthorrhiza, Pistia stratiotes*, *Salvinia rotundifolia;* Fig. S1) and high occurrence of littoral and benthic diatoms (e.g. *Eunotia, Encyonema* and *Pinnularia;* Fig. S3b) and charophyte macro algae (*Chara* spp. and *Nitella* spp. Fig. S1), all of which commonly occur in low turbulence and low nutrient waters^37–39, 41^. The occurrence of planktonic diatoms (e.g. *Aulacoseira* spp., Fig. S3a) further indicates an environment hydrologically connected to a main river channel^37–39^. Abundant Trichoptera shredders and macrophyte/detrital-associated invertebrates (e.g. *Cladopelma* spp*., Zavreliella* spp., and *Stenochironomus* spp.; Fig. S2) along with anaerobic bacteria-archaea all suggest a highly reductive environment. In particular, the latter suggests that carbon cycling at the time might have mainly occurred through methanogenesis and sulphate reduction pathways^42^.

**Figure 4.**
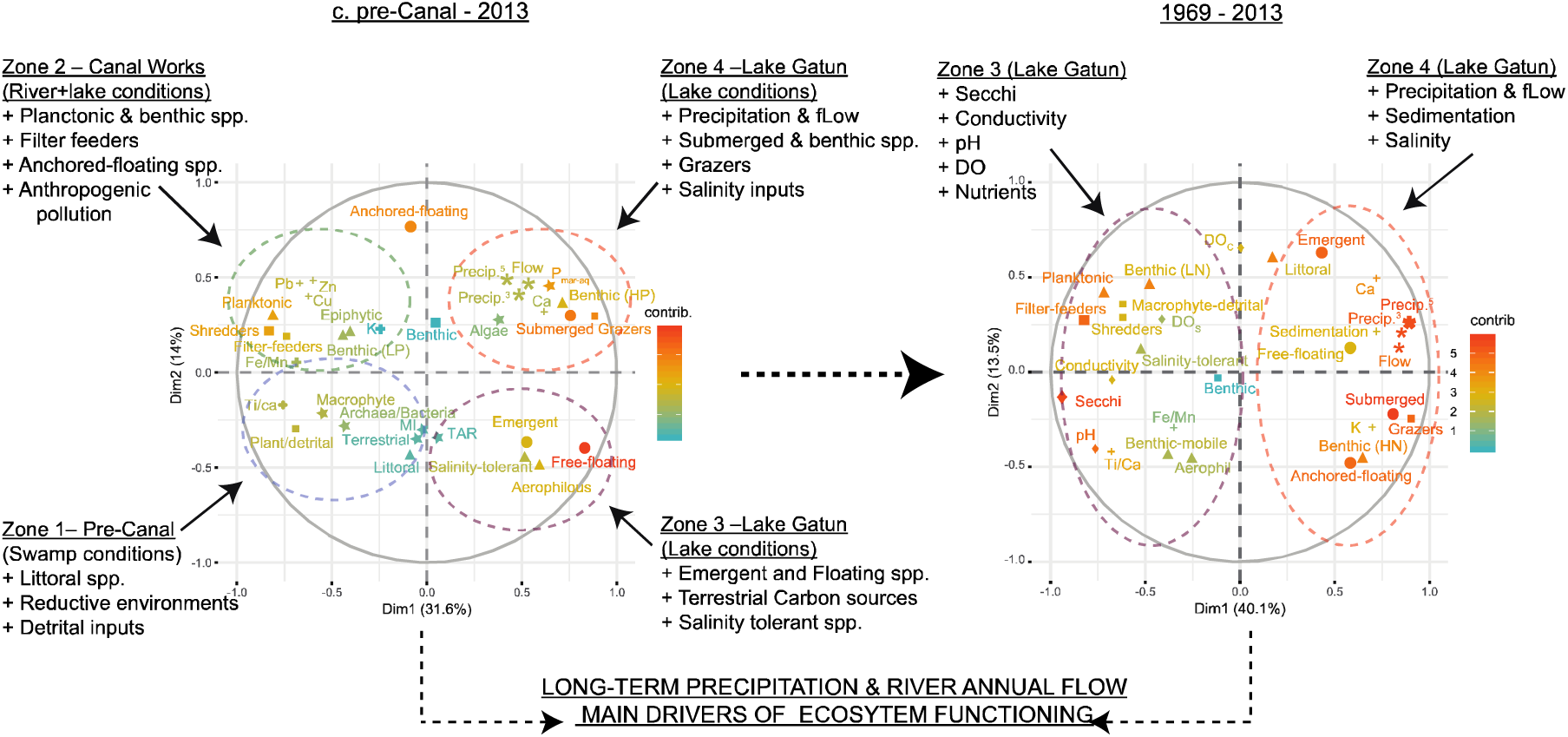
Multiple factor analysis (MFA) results for the time periods *c.* pre-canal-2013 and 1969-2013. The analysis was run on hydro-climatic data (precipitation and river flow–hash), biological data (macrophytes–circle, diatoms– triangle, invertebrates–square), geochemical data (cross), biomarker data (star) and water quality data (period 1969-2013–diamond). The contribution of each variable in the analysis is indicated according to a color scale. Low nutrients (LN), high nutrients (HN), three years average precipitation data (Precip. ^3^), five ars average precipitation data (Precip.^5^), river annual flow (Flow). Sediments samples within each major temporal zone of compositional change are encircled according to Zone 1 c. pre-1870, Zone 2 c.1870-1914, Zone 3 1923-1990, and Zone 4 1991-2013.

Construction of the Panama Canal led to a clear anthropogenic signal (Fig.2). From 1870-1913 (Zone 2) pollutants such as Cu, Zn, and Pb, previously associated with mining activity, coal burning and gasoline combustion^43^, increased considerably. These activities were most likely linked to the coal combustion by large machinery used for the dredging and excavation of the canal^40^. Submerged (e.g. *Najas guadalupensis, Najas marina*, and *Ceratophyllum demersum;* Fig. S1) and anchored-floating plants (likely *Nymphaea ampla*), benthic-mobile diatoms (e.g. *Navicula radiosa* and *Navicula recens;* Fig. S3c), caddisfly larvae (Trichoptera) shredders and bryozoan filter-feeders (*L. carteri*, and *Plumatella* spp.) responded positively to these novel conditions while diatoms shifted from littoral to benthic-planktonic associations (Fig. 4). Increases in the bryozoans *L. carteri* and the colonizing *A. gelatinosa* in particular, were likely responding to an expansion of macrophyte cover^26, 44^. The observed proliferation of caddisfly larvae may also have ultimately been driven by increased detrital inputs and food availability as they prey on bryozoans^44^.

As the lake infilled after formation of the dam in 1913 (Zone 3), littoral detrital (Ti/Ca) and river-fed elements (e.g. K) declined (Fig. 2). Declining erosion after dam construction likely continued following construction of the Alajuela Dam in 1935 in the headwaters of the Chagres, reducing the supply of river material into Gatun Lake^15^. However, contributions of allochthonous organic carbon (high TAR index) increased up until the mid-1980s (Fig. 2). Possible sources of this terrestrial carbon include the flooded forest areas^13^ and particulate material derived from watershed deforestation, which peaked during the mid-1970s^16^. These patterns of increasing terrestrial organic matter inputs from early-1920s to the mid-1980s partially supports recent findings showing that the degradation of flooded forest material in tropical impoundment projects may endure for decades after reservoir infilling, a period when CO_2_ and CH_4_ production is commonly facilitated^7, 18^. Yet, our data suggest that such carbon pathways may take even longer to develop (four-five decades) than previously suggested for tropical dam projects^7, 18^.

In general, our findings show that macrophyte growth was encouraged as the lake infilled, promoting aerobic conditions (Fig. 4). This trend matches the historical macrophyte records from Gatun Lake^24^, and other similar tropical impoundment projects^8, 45^. For instance, the invasion of *H. verticillata* that resulted in many hectares of the lake becoming choked with this submerged species was accompanied by increases in other submerged plants like *Najas* and *Ceratophyllum* (Fi. S1). Free-floating plants, such as *Eichhornia spp.*, and *P. stratiotes*, also disseminated rapidly, while anchored plants (*N. alba* in particular), invertebrate shredders, filter-feeders and macrophyte/detrital associated taxa declined (Fig. 3, Figs. S1,2.).

Diatom and invertebrate assemblages mirrored the trends in macrophytes following impoundment (Fig. 4). Over time, invertebrate communities shifted from detrital to benthic associations, while bryozoan filter-feeders declined (Fig. 4), which could have been a response to the gradual decline of anchored-floating plants^26, 44^. Diatoms shift to a benthic-aerophilous-saline diatom assemblage characterized by increases *Cocconeis placentula* and *N. amphibia*, by aerophilous genera such as *Diadesmis, Luticola* and *Orthoseira* and by the salinity-tolerant species *Terpsinoe musica*, and *Tabularia fasciculate*^35–37^(Fig. S3). This shifts in diatom composition suggest that the progressive macrophyte expansion provided an increase in habitat availability for benthic-mobile species and suitable macrophyte littoral habitats for aerophilous species^38, 39^ while increases in saline-tolerant taxa were most likely driven by salt-water intrusions from the passage of ships through the dam locks^20^ and ion runoff resulting from deforestation^16^.

As the lake aged (post-1995; Zone 4) submerged and free-floating macrophytes increased and carbon cycling shifted in concordance to within-lake production (high Pmar-aq index value)^31^ (Fig. 2, 3). This shift in habitat structure marked an upsurge in the abundance of grazing invertebrates and benthic diatom species that prefer productive environments (fig. 2).

### Impoundment and natural river floodplain dynamics

We found that after a century of the dam construction, there are still remarkable parallels associated with the natural hydrological functioning of a river floodplain system. In particular, precipitation and annual river flow emerged as the most important factor driving most of both abiotic and biotic compartments (Fig. 4) in agreement with floods and droughts being major drivers of river abiotic change and community reorganization^46^. Shifts in hydro-climatic variables were suggested to alter a series of interconnected processes such as sedimentation dynamics, water quality, productivity, and sediment reductive/oxygenated conditions. During drier periods for instance, sedimentation was relatively low due to lower physical erosion while detrital inputs where high (Fig. 4). There was also a prevalence of reductive sediments and relatively higher secchi depths (> 3m), oxygenated surface waters (> 6 ppm), higher conductance (>60 μS/cm) and higher nutrient availability (Table 1; Fig. 4). Similar increases in conductance resulting from reduced dilution of salt ions concentrations during the dry season have been observed in the Amazon River, where conductance values in oxbow lakes can increase up to 200 times the value of the main river^46^. Accumulation of organic matter and debris in the lakebed causing reductive soil conditions have been also described in the Paraná River system and attributable to low rates of water circulation during drier phases^47^. The observed anoxic sediment conditions (high Fe/Mn ratio) in Gatun, would have transformed nutrients (phosphorus in particular) into more available forms^42^ that along with clearer and less variable water levels would have favored planktonic diatoms^37–39^ and submerged and free floating macrophytes^45, 47^. As submerged plants grow in clearer waters they would have also photosynthesized more increasing surface DO levels in the water^48^.

**Table 1.** Historical environmental records of Lake Gatun. Historical data on three years average precipitation data (Precip.^3^), five years average precipitation data (Precip.^5^), and annual river flow in Gatun Lake for the periods 1930-2013, and historical data on nitrates (NO_3_), phosphates (PO_4_), secchi depth, conductivity, pH, dissolved oxygen at the water surface (DO_s_) and at the water column (DO_c_ < 1m depth), and chlorophyll a (Chl-a) for the period 1969-2013.

| Time<br>(yrs.) | Precip.<br><sub>5</sub><br>(mm) | Precip.<br><sub>3</sub><br>(mm) | Annual<br>flow<br>(m3/s) | pH | DO<sub>s</sub><br>(mg/L) | DO<sub>c</sub><br>(mg/L) | NO<sub>3</sub><br>(mg/L) | PO<sub>4</sub><br>(mg/L) | Cond.<br>(μS/cm<br>) | Chl-a<br>(μl/L) | Secchi<br>(cm) |
| --- | --- | --- | --- | --- | --- | --- | --- | --- | --- | --- | --- |
| 2013† | 2964 | 3381 | 41 | 6.41 | 6.44 | 5.25 | 0.04 | 0.01 | 42.34 | 2.31 | 146 |
| 2011† | 2841 | 2778 | 38 | 6.35 | 6.60 | 5.12 | 0.07 | 0.01 | 47.86 | 3.22 | 150 |
| 2009† | 2664 | 2640 | 31 | 6.50 | 6.48 | 5.01 | 0.03 | 0.00 | 49.38 | 4.18 | 167 |
| 2008† | 2769 | 2741 | 31 | 6.59 | 6.28 | 4.23 | 0.03 | 0.01 | 55.62 | 5.09 | 177 |
| 2006† | 2569 | 2685 | 29 | 6.56 | 6.25 | 3.51 | 0.04 | 0.02 | 55.62 | 2.89 | 212 |
| 2005† | 2620 | 2689 | 27 | 6.53 | 6.23 | 2.52 | 0.03 | 0.01 | 45.21 | 4.72 | 192 |
| 2003† | 2593 | 2802 | 30 | 7.13 | 6.61 | 2.90 | 0.03 | 0.02 | 50.33 |  |  |
| 2000† | 2688 | 2644 | 37 | 6.76 | 6.48 | 3.02 | 0.05 |  | 49.92 |  |  |
| 1993§ | 2589 | 2669 | 28 | 7.39 | 6.22 | 1.94 |  | 0.04 | 54.00 |  |  |
| 1989§ | 2517 | 2524 | 26 |  | 8.66 | 4.94 | 0.03 | 0.04 | 44.88 | 2.05 | 393 |
| 1978* | 2508 | 2332 | 25 | 7.20 | 7.78 |  | 0.06 |  | 90.00 | 4.10 | 700 |
| 1972* | 2331 | 2371 | 26 | 7.56 | 8.00 |  | 0.05 | 0.02 | 98.00 |  |  |
| 1969* | 2497 | 2444 | 27 | 7.21 | 5.59 | 5.12 |  |  |  |  | 530 |
| 1955 | 2663 | 2859 | 29 |  |  |  |  |  |  |  |  |
| 1930 | 2679 | 2488 | 27 |  |  |  |  |  |  |  |  |

\*ref-53; §ref-56; †ref-57; precipitation data were obtained from (Steve Paton, pers. comm.) and annual flow data from ref-23.

Wet periods positively stimulated sedimentation, deposition of new minerals (e.g. Ca and K) and reduced water transparency (Fig. 4). Lower secchi depths were strongly linked to reductions in pH, conductance and dissolved oxygen (Fig. 4), which likely reflects the storage of organic matter during the dry phase, coming from both autochthonous production and allochthonous inputs from the lavish riparian vegetation surrounding the lake catchment^47^. A positive long-term relationship between macrophyte productivity and wet periods also existed (Fig. 4). We attribute this increase in productivity to a decadal macrophytes succession that was accompanied by increases in abundances and representation of different plant taxonomic groups (Fig. S1). Furthermore, high rates of river water exchange during high floods can act as a significant source of propagules, especially for submerged and floating plants in neotropical rivers^47^. The action of water transporting sediments and nutrient (TP and TN) additions from surrounding soils, may promote further spatial heterogeneity in the lake^46, 47^ opening up additional ecological niches for macrophytes and co-associated benthic diatoms. Light attenuation in the water column coupled with fluctuation in water levels may further stress the submerged vegetation while favoring increases in floating macrophytes through enhanced nutrient input from the flooded land^45, 47^. The weak negative relationship between macrophyte abundance and nutrient concentration (Fig. 4), suggests that primary productivity in the lake may not yet be limited by nutrients. Moreover, changes in habitat structures (anchored floating plants) rather than nutrient availability are suggested to favor the variation of nutrient-rich associated benthic diatoms.

We are aware that the use of paleoecological data to infer past communities and ecological responses has limitations as they can suffer from bias due to the differential production, transport and preservation of organismal remains^49^. Nevertheless, our paleoecological data was in agreement with the historical changes previously described for the lake. Uncertainties in the age model and use of different historical environmental records might have also introduced some discrepancies in our MFA model. Nevertheless, the observed limnological changes agree with the literature of river floodplain dynamics^46, 47^. This thesis is further supported by the geochemical data that showed a coherent signal with increases in precipitation over 2010-2011. We observed a drastic decline in Ca coupled with increases in sedimentation rates, Fe/Mn, and Ti/Ca (Fig. 2), which resemble the historical riverine pre-damming conditions and that are in agreement with “La Purísima” rainstorm, which flooded the whole lake system and increased sedimentation rates by almost 100-fold^50^.

### Is the Gatun Lake becoming more saline?

Our data are concordant with a gradual increase in salinity in some parts of Gatun Lake after the early-2000s as observed by increases in saline-tolerant diatom species and Ca concentrations. Seawater likely intrudes into the lake through the locks of the Panama Canal and the deposit of ballast water into the lake, which was only forbidden after 1996. The locks may not be the only reason for increasing salinity. Runoff associated with the enclosed drainage basin and land-use change may have led to an increase in ion input and hence increased water salinity. These two processes are often governed not only by greater runoff but also by evaporation; a pattern that is consistent with the peak in salinity-tolerant diatom species and increases in emergent macrophytes during the drier period of the mid-1960s to mid-1980s (Figs. 3, 5b). The more recent increases in salinity-tolerant diatom species (including two marine morphtypes; Fig. S3f) in the Gatun Lake after the early-2000s implies an increase in the rate of salinization of the lake. This coincides with the canal expansion work, which began in 2005 and the use of new locks built to permit the transport of larger vessels (Post-Panamax) through the canal^20^. These new locks use a tiered water sharing system that can more easily move salt water up into the lake easier. Recent STRI salinity monitoring data supports this inference (Steve Paton, pers. comm.). As deforestation in the lake catchment has declined in the last two decades^17, 24^, it is likely that runoff patterns might not have been playing such an important role in salt intrusions in comparison to the locks.

If the predicted drying of the canal area due to global climate change is correct^12^, and global shipping traffic increases^28^, it is likely that salinity will continue to increase in the lake with major implications for drinking water^17^. The ecological consequences of increased salinity has yet to be properly explored but the recent increase we observe in *n*-alkane bacteria and both the MI and Pmar-aq indices could be a warning that the halocline will render surface sediments anoxic^51^.

### Ecological responses to fish invasion

The invasion of the apex predator *Cichla* (peacock bass) into Gatun Lake in 1969 had a profound impact on the native littoral planktonic fish community^25^, which resonate today with marginalized native populations^23^. Zaret and Paine^24^ predicted that predation by *Cichla* would also lead to cascading effects through the lake’s food web, particularly on zooplankton (e.g. *Ceriodaphnia*), aquatic insects (e.g. mosquitos/chironomids) and primary producers. Our results however do not support the latter, as cladoceran ephippia only became apparent after the late 1990s, a period that instead coincides with increasing *n*-alkane algae contribution. We found no evidence of increasing abundances or shifts in specific functional groups (e.g. planktonic taxa) during or post-*Cichla* times that would support evidence for a long-term cascading effect of predation down the food web. Instead, evidence during this period points towards a bottom-up flow of energy and nutrients coupled with asymmetric benthic–littoral production likely associated with the development of macrophytes^52^.

### Remarks and management options

Our data help reconstruct the biotic and abiotic dynamics of the Chagres River landscape over the last ∼150 years, providing a unique insight into the natural and anthropogenic impacts of impoundment on tropical rivers. Species invasions, land-use changes and ship traffic have impacted the lake’s ecosystem. Yet, our multiple lines of evidence emphasize that the system still retains some of the natural riverine functions on a decadal scale. It is however anticipated that climate change will modify precipitation, evapotranspiration, and runoff in the tropics^12^. Thus, increasing drier and wetter periods could fundamentally modify the functioning of Gatun Lake. Drier periods will likely encourage the on-going spread of *Hydrilla* and *Eichhornia* and increases in salinity via reduced dilution. Wetter periods in turn, may stimulate sedimentation rates, nutrient inputs, salt intrusions from storm surges and floating plant dominance. Many neotropical impoundment projects have become highly eutrophic within a few decades, rapidly overriding the importance of hydrology^8, 10, 45^. However, Gatun Lake stands apart, by having high precipitation rates (annual mean > 2.200 mm), a unique occurrence of large extensions of protected forest areas (e.g. Barro Colorado Natural Monument and Soberanía) in the lake catchment, and large quantities of water leaving the system (> 1 million m^3^/year)^53^ every time a ship pass through the lake locks. These factors may provide some buffering, helping to reduce shifts in runoff, water pollution and maintaining the natural hydrological balance under a changing climate. Our study emphasizes that to preserve natural riverine system functioning in tropical impoundment projects, management activities must not only include better design and management of flow releases^3^ but also the understanding of key long-term natural ecosystem structural drivers such as river flow, runoff patterns and water exchange rates.

## Materials and Methods

### Study area

Gatun Lake is situated in the valley of the Chagres River to the south of Colón, Panama (9°11′N 79° 53′W) (Fig. 1). It is an artificial large lake (425 km^2^) with a maximum water depth of 30 m and extensive areas of shallow water (<5 m). The lake waters are well mixed throughout much of the dry (mid-December to mid-April) and wet (mid-April to mid-December) season^53^. The lake level is 26 m above sea level storing 5.2 km^3^ of water^53^. It serves a dual purpose, as a channel facilitating global trade and cross-oceanic travel, and as a freshwater reservoir (Gatun Lake) providing a water supply to Panamá City and other towns^17^.

### Sampling

A sediment core (87 cm long; LGAT1) was retrieved in 2013 from near “La Represa” village in the southeast area of the lake (Fig. 1). The basin offered an ideal coring site given that it lies outside the dredging zone of the canal and is located in one of the most deforested areas of the lake watershed. The core (LGAT1) was retrieved from a littoral area (9° 2’49.58″N, 79°50’6.33″W; Fig. 1) at a water depth of approx. 1 m. We used a modified Livingstone Piston Sampler of 4 cm diameter. Sediment samples were extruded in the field at 1 cm intervals.

### Core dating

Fourteen dried sediment samples from core LGAT1 were analyzed for ^210^Pb, ^226^Ra, ^137^Cs and ^241^Am by direct gamma assay in the Environmental Radiometric Facility at University College London, using ORTEC HPGe GWL series well-type coaxial low background intrinsic germanium detector. The sedimentary chronology was determined using the Constant Rate of Supply model (CRS)^34^.

### Geochemical elements

Elemental composition on 1cm-thick discrete samples was measured via X-Ray Fluorescence (XRF) on a handheld XRF analyzer spectrometer, XMET 7500. Dry sediment samples were ground and homogenized using a mortar and pestle. Three g of sediment sample were used and covered with a Chemplex thin-film sample support. The handheld XRF analyzer spectrometer was calibrated against certified material prior to analysis and mean values for each element were determined from duplicate measurements. Sampling resolution was at 2-cm intervals for the top 50 cm of the core and at 4-cm for the remainder. A total of 34 samples were analyzed. Calcium (Ca), potassium (K), iron (Fe), manganese (Mn), titanium (Ti), lead (Pb), copper (Cu) and zinc (Zn) data were selected for this study. The elements Pb, Cu and Zn were used as proxies for human-derived pollution events, Ca as a proxy of marine influence, and K as a proxy of physical erosion^56^. We calculated complementary index ratios to investigate changes in reduction conditions (Fe/Mn) and detrital input (Ti/Ca.)^29^.

### Biomarkers

We analyzed *n*-alkanes composition in 10 sediment samples^30^. We used *n*-alkanes C_15_–C_31_, as indicators of terrestrial plants, macrophytes, and bacteria-algae^30^. Compounds were measured with a Shimazu GC-2010 gas chromatograph interfaced to a Shimazu GCMS-QP2010 (see ref–58 for detailed methodology).

Besides relative *n*-alkane contribution, we also calculated the terrigenous aquatic ratio (TAR), which quantifies the *in situ* algal vs. terrestrial organic matter^31^ and the submerged/floating aquatic macrophyte inputs *vs*. emergent/ terrestrial plant input ratio (Pmar-aq)^31^. The Pmar-aq quantifies the non-emergent aquatic macrophyte input to lake sediments relative to that from the emergent aquatic and terrestrial plants^31^. Values of Pmar-aq <0.1 corresponds to terrestrial plants, of 0.1-0.5 to emergent macrophytes and of >0.5-1 to submerged/floating macrophytes^31^.

The methane index (MI) that quantifies the relative contribution of methanotrophic Euryarchaeota against ammonia oxidizing Thaumarchaeota was also calculated^32^. MI values close to 1.0 indicate anaerobic environments, whereas values close to zero indicate aerobic conditions^32^. To calculate the MI index we measured glycerol dialkyl glycerol tetraethers (GDGTs) using an Agilent 1260 UHPLC coupled to a 6130 quadrupole MSD high performance liquid chromatography-atmospheric pressure chemical ionization-mass spectrometry (HPLC-APCI-MS).

### Plant and invertebrate macrofossil

We analyzed 23 sediment samples for plant and invertebrate remains. Between 2-4 g of dried sediment material per sample were used and all samples were disaggregated in 10% potassium hydroxide (KOH) before sieving. Macrophyte fossils were retrieved from the residues of sieved core material (using mesh sizes of 355 µm and 125 µm) following standard methods^54^. Plant macrofossil data were standardized as the number of fossils per 100 cm^3^ and identified by comparison with reference material and by using relevant taxonomic keys^54^. Due to poor preservation of *Hydrilla* remains, we estimated temporal abundances through its well-documented historical records^23^ and expressed in a 0-3 scale, where 0 is absent and 3 highly abundant. All macrophyte taxa were then classed based on preferred growth-type as submerged; anchored-floating; free-floating and emergent. Invertebrate taxa were classed according to feeding behavior or preferred habitat as: filter-feeders (bryozoans); macrophyte/detritus (chironomids), shredders (Trichoptera larvae); benthic (chironomids), and grazers (mollusks and cladocerans).

### Diatoms

Twenty-three samples were analyzed for diatoms following standard procedures by Battarbee et al.^55^. The diatom suspension was mounted on slides with Naphrax® after the removal of carbonates by HCl and organic matter by H_2_O_2_. Diatom taxonomy and ecology mainly followed Diatoms of North America (https://diatoms.org). Diatom species were classed based on preferred habitat type as planktonic, benthic, littoral and aerophilous, and as salinity-tolerant. Benthic species were sub grouped into acidic-oligotrophic, eutrophic, and mobile species. For each sediment sample, we counted a minimum of 400 valves.

### Historical environmental archives

Historical data on water quality and hydro-climatic variables are presented in Table 1. Long-term hydro-climatic, i.e. precipitation (three and five years average) and river annual discharge data, from 1930 to 2013 were obtained from the monitoring division of STRI (Steve Paton, pers. comm.) and the Panama Canal Authority (23). Long-term physical and chemical variables from 1969-2013 (pH, conductivity, dissolved oxygen [DO], total nitrogen [TN], total phosphorous [TP], chlorophyll a [Chl-a] and secchi depth) were obtained from literature^56, 57^ and from the ACP Water Quality Monitoring Division reports^57^. For the ACP data we used the average annual values of each selected variable recorded at two sampling stations (Laguna Alta– LAT and Toma de Agua Represa–TAR) located near our coring site area.

### Data analysis

A stratigraphic plots of the study biological functional groups were achieved using the ‘‘Rioja’’ Package in R^58^. Major zones of change were achieved through clustering analysis. To summarize and visualize the most important gradients of temporal change in the different biological functional groups, environmental variables, and historical data we used multiple factor analysis (MFA)^33^ in R (*Factoextra* and *FactoMiner* packages in R). This analysis is a multivariate technique in which individuals (core depths) are described by several sets of variables (quantitative in our case) structured into groups. The analysis takes into account the contribution of all groups of variables to define the distance between core depth samples. Variables within a group were normalized by applying a weight equal to the inverse of the first eigenvalue of the analysis of the group^33^.

We ran two separate MFA analyses according to data availability. The first analysis focused on the historical hydro-climatic variables (river flow and precipitation; n=15 data points) and all paleoecological data spanning since pre-canal times and clustered into five major groups: macrophytes (submerged, anchored-floating, free-floating and emergent; n=23 data points for each group), diatoms (littoral, benthic, planktonic, salinity-tolerant, and aerophil; n=23 data points for each group), invertebrates (filter-feeders, shredders, benthic, macrophyte/detritus and grazers; n=23 data points for each group), geochemical elements and ratios (n=23 data points for each element or ratio), and biomarkers; (n=10 data points for each parameter). Missing information at given time-periods for biomarkers (10 samples analyzed) and hydro-climatic data (spanning from 1930-2013), was replaced in the MFA by the mean of each variable^33^. The second MFA focused on the time-period 1969-2013 from which historical water quality data (pH, conductivity, phosphorous, chlorophyll-a, and secchi depths; n=13 data points) was available (Table 1). For this analysis we included the historical hydro-climatic data (n=13 data points) and the paleoecological groups data (n=13 data points for each biological and geoechemical group), excepting biomarkers (too limited number of samples), from analysis 1. To quantify the contribution of each variable in the MFA maps we used the argument col.var = “contrib” (*Factoextra* package, R). Prior to MFAs we run exploratory principal component analyses (PCA) to detect uninformative or redundant historical water quality variables (see Fig. S4 for details). As a first step we discarded nitrates due to low variation (Fig. S4b) and subsequently, we discarded phosphates and chl-a as these two variables showed a strong correlation with DO at the water surface (positive) and with DO in the water column (negative) respectively (Fig. S4c). We left both DO measurements instead of nitrates and chl-a due to a better historical record (see Table 1).

### Data Availability

all relevant data are available from the authors and the data will be deposit in a public repository.

## Acknowledgments

We thank the Smithsonian Tropical Research Institute (STRI) for funding fieldwork and supporting JS through a Short-term Postdoc Fellowship. We thank Dr. Dolores Piperno at STRI, for lending us the coring device. We thank Universidad de Los Andes and COLCIENCIAS for supporting JS under the postdoctoral program ‘‘Es tiempo de volver’’ Convocatoria 2015. We also thank Vicerrectoria de Investigaciones of Universidad de Los Andes, for supporting one month of salary for JS. We thank the Bloomsbury Environmental Isotope Facility at University College London for sediment dating. We thank Victor Frankel, Felix Rodriguez, Luis J. de Gracia, Marcos Alvarez, Maria Pinzon and Marcela Herrera for fieldwork, laboratory assistance and hospitality. We thank the Geociencias Laboratory at Universidad de los Andes for facilitating the XRF analyzer and Laura Caceres for analyzing the XRF data. We thank Steve Paton for historical climatic data provision. We want to thank Professors J.-H. Kim and K.-H. Shin for the measurement of the biomarkers in their laboratory at the University of Hanyang, South Korea. The collection and exporting of sediment material was assessed under the ARAP collecting permit No. 25.

## Author Contributions

JS designed the study and collected the sediment material. JS produced and analyzed the plant and invertebrate macrofossil data, CH and JC produced and analyzed the biomarker data, and MV produced and analyzed the diatom data. NR and HY produced and analyzed the lithostratigraphic and radiometric data. JS wrote the first manuscript and all authors contributed substantially for the final version.

