## Supplementary data for "The Panama Canal after a century of human impacts"

*corresponding author: Jorge Salgado

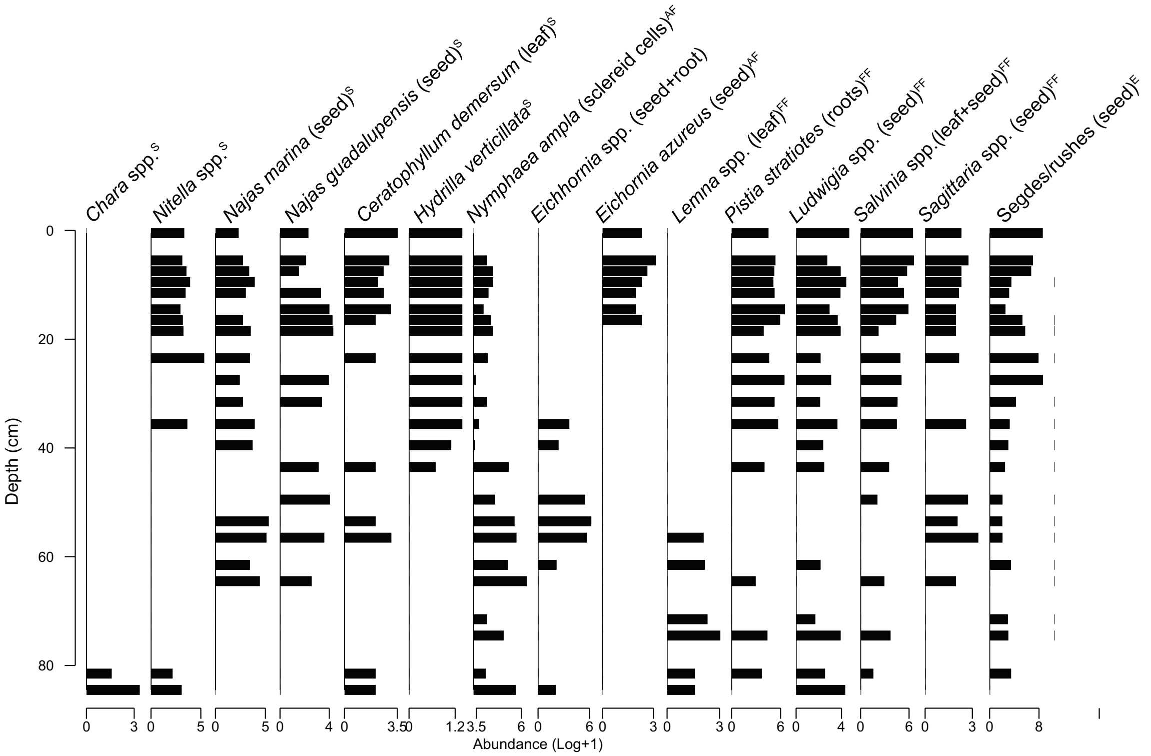


**Figure S1**. **Sedimentary profile of plant macrofossil taxa in LGAT1 sedimentary core.** Growth type of each taxa is indicated in superscript as: submerged (S), anchored-floating (AF), free-floating (FF), and emergent (E).

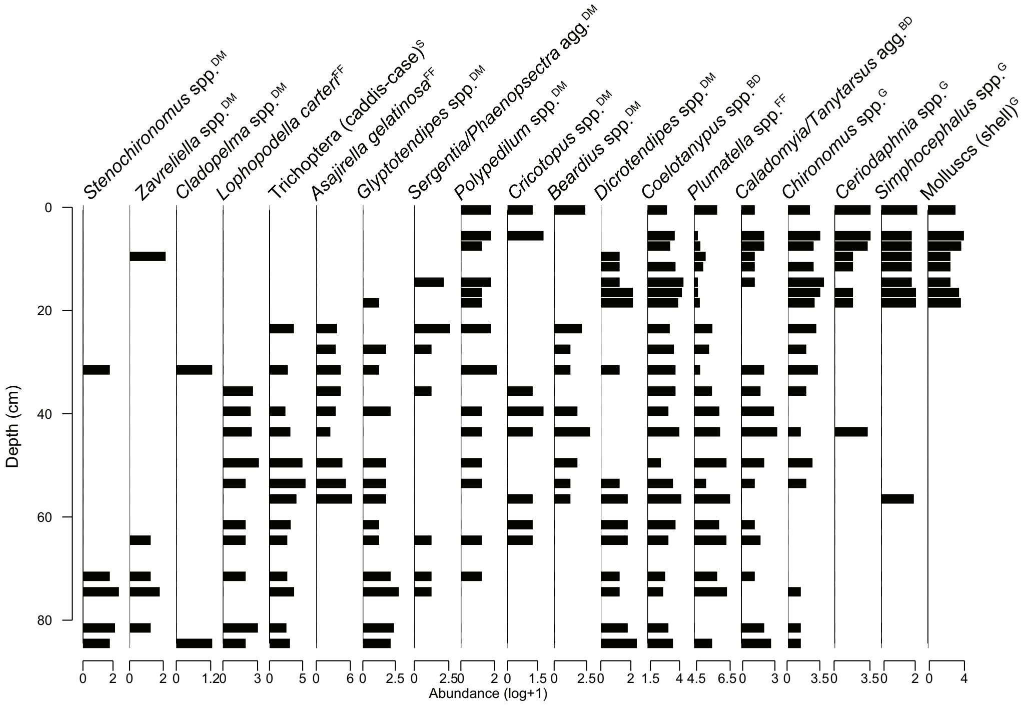


**Figure S2**. **Sedimentary profile of invertebrate taxa in LGAT1 sedimentary core**. Selected feeding behavior or preferred habitat of each taxa is indicated in superscript as: filter-feeders (FF), grazers (G), benthic-dam (BD), shredders (S), detrital-macrophyte (MD).

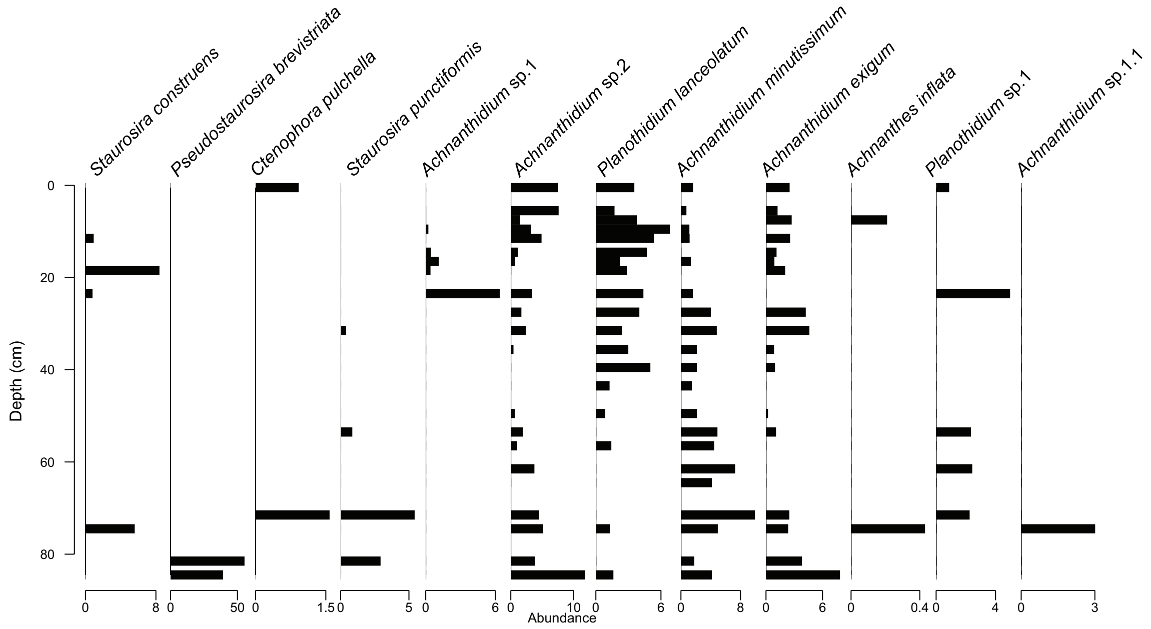


**Figure S3a**. **Sedimentary profile of littoral diatom species in LGAT1 sedimentary core.**

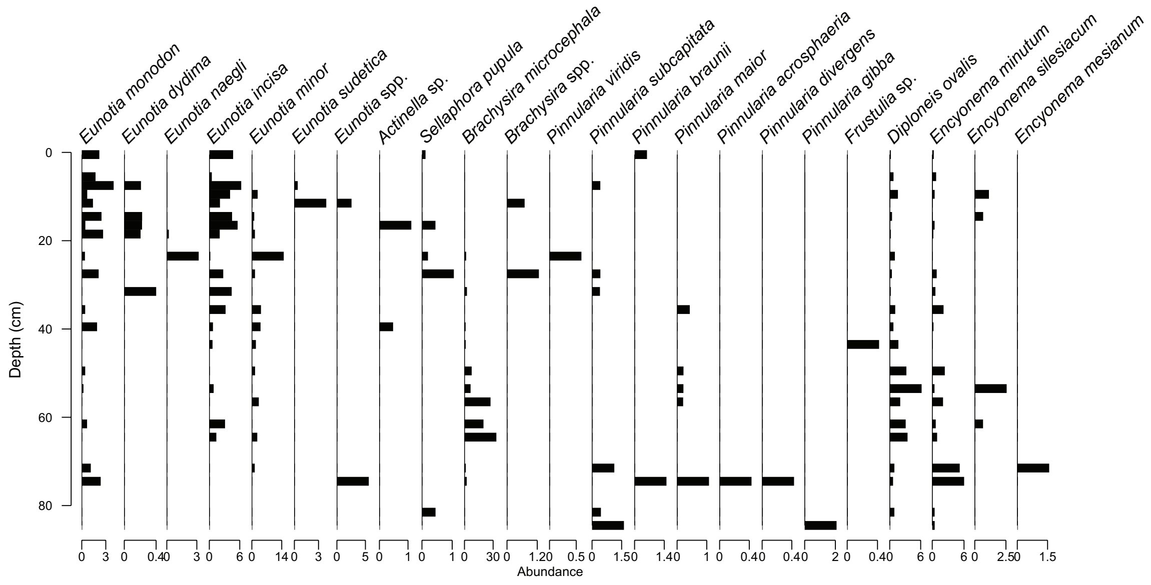


**Figure S3b**. **Sedimentary profile of benthic (low nutrients) diatom species in LGAT1 sedimentary core.**

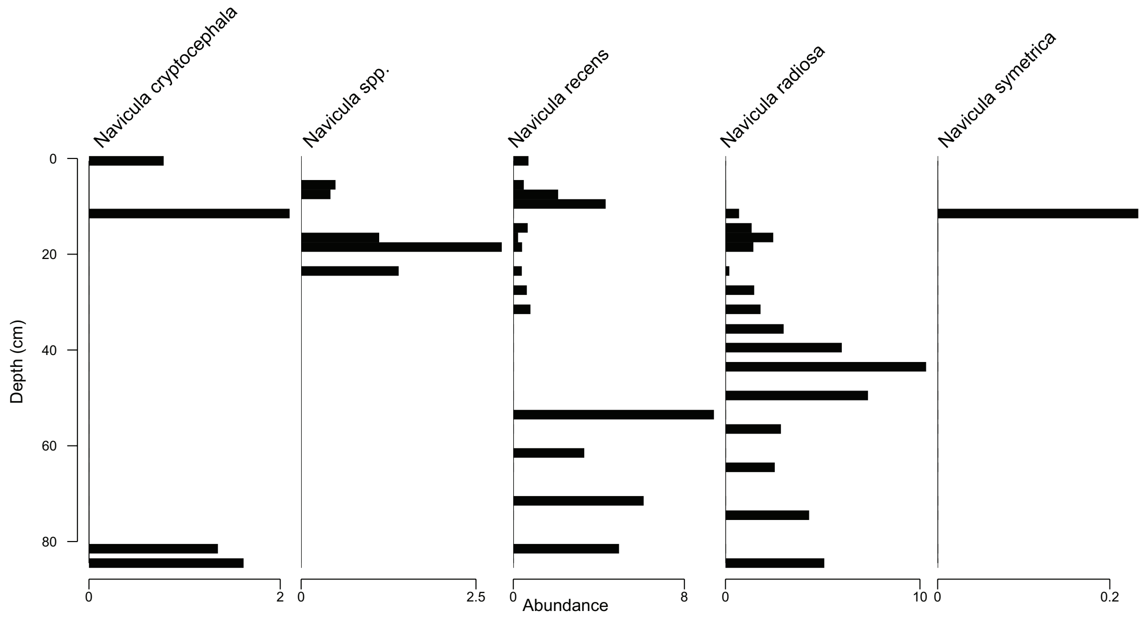


**Figure S3c**. **Sedimentary profile of benthic-mobile diatom species in LGAT1 sedimentary core**.

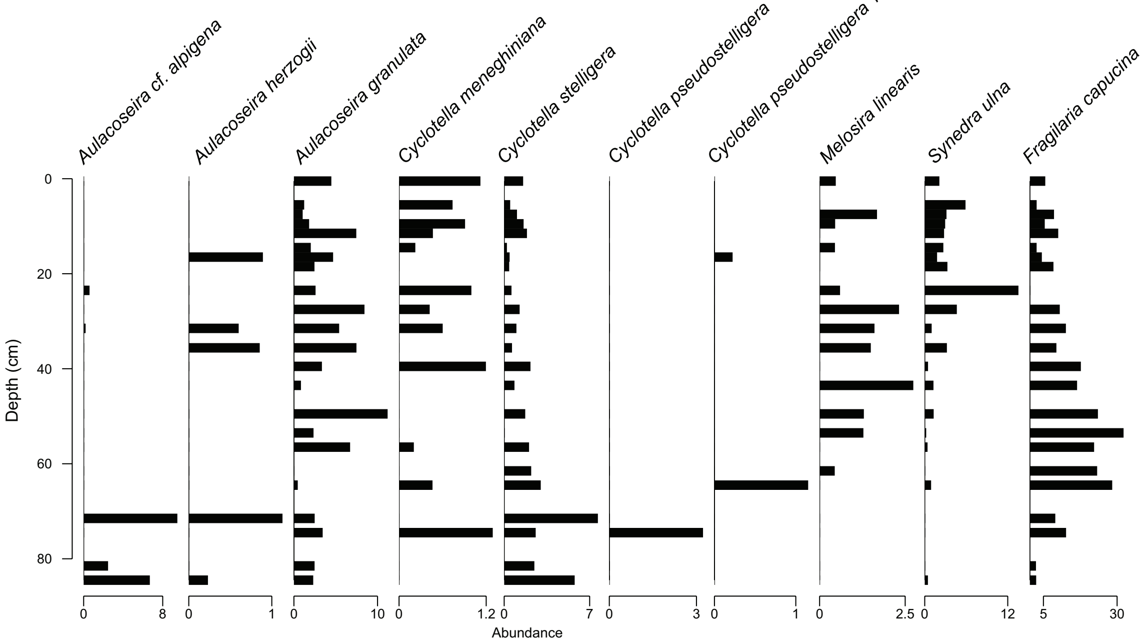


**Figure S3d. Sedimentary profile of planktonic diatom species in LGAT1 sedimentary core.**

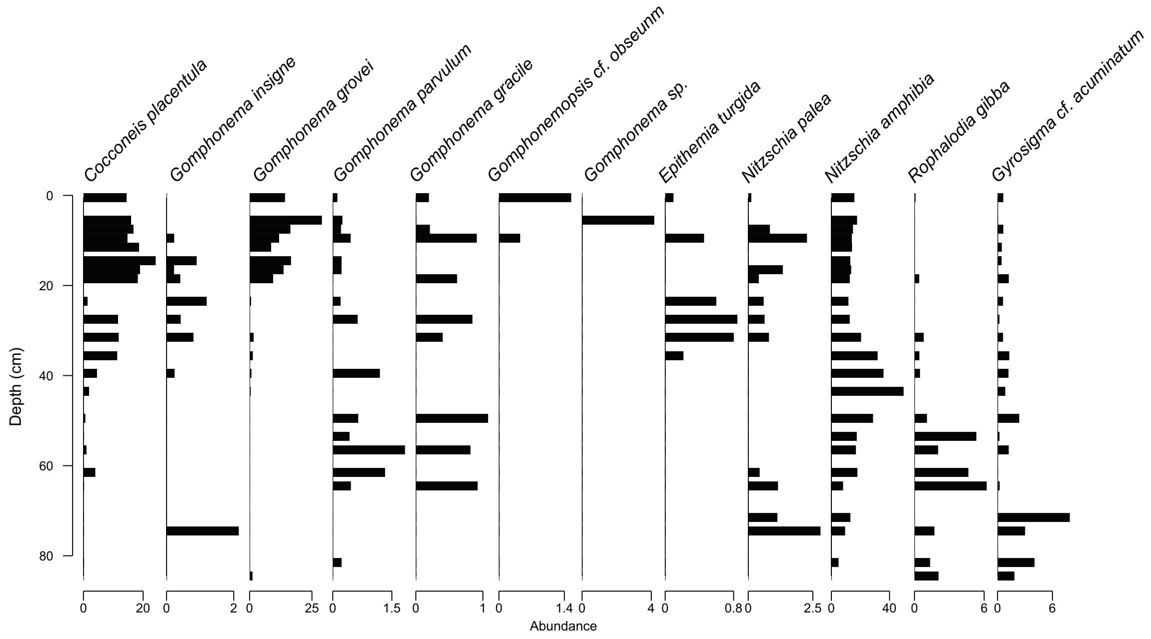


**Figure S3e**. **Sedimentary profile of benthic-rich diatom species in LGAT1 sedimentary core.**

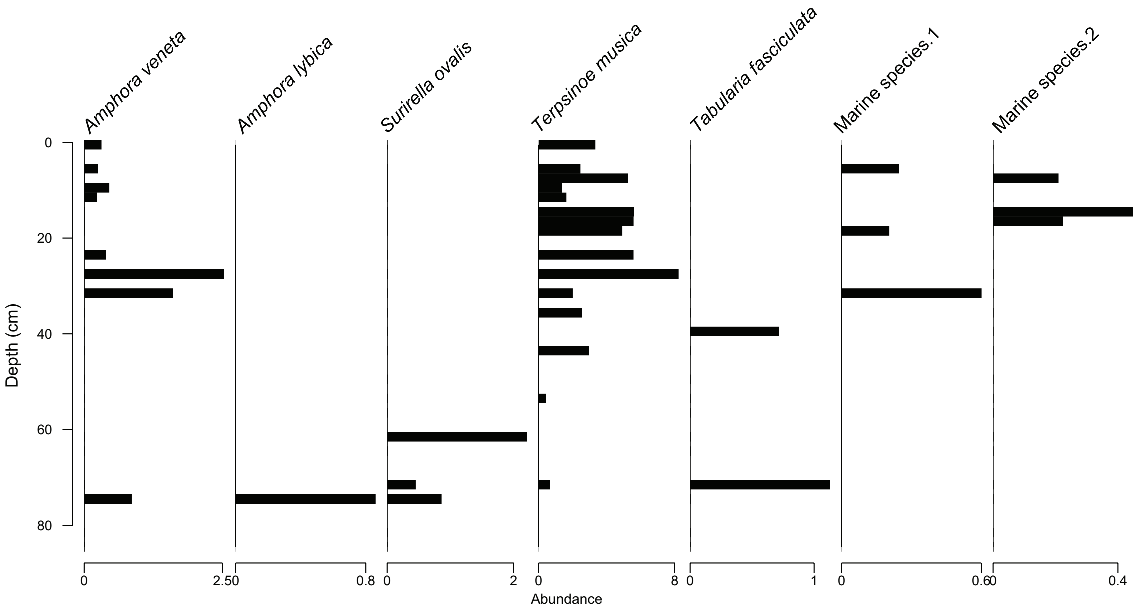


**Figure S3f. Sedimentary profile of salinity-tolerant diatom species in LGAT1 sedimentary core.**

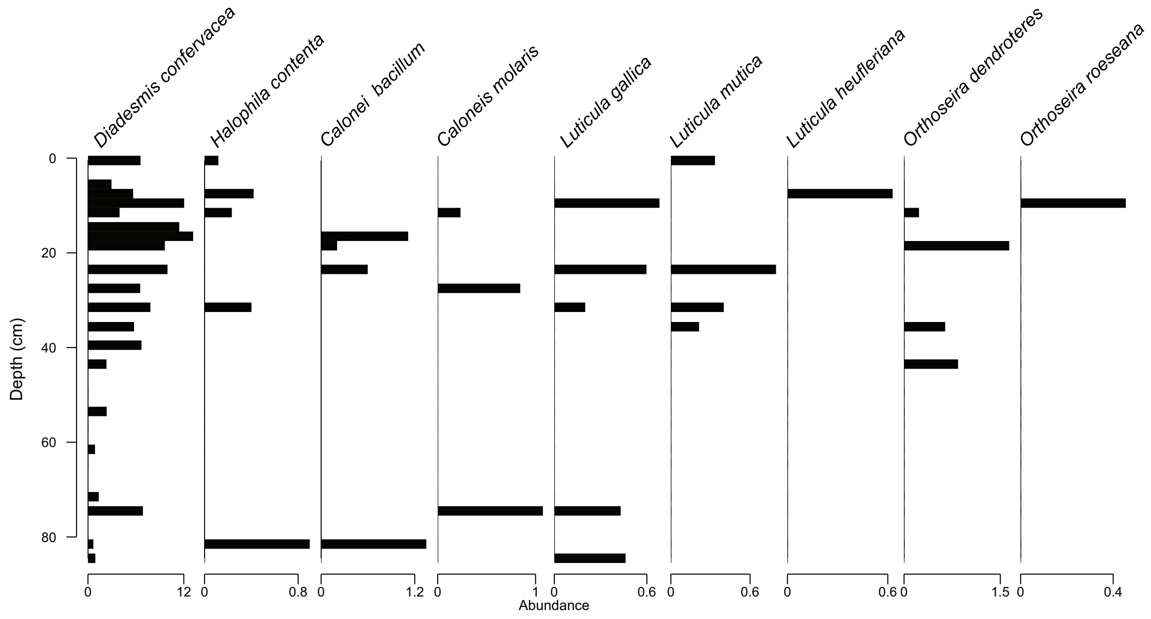


**Figure S3g**. **Sedimentary profile of aerophilous diatom species in LGAT1 sedimentary core.**

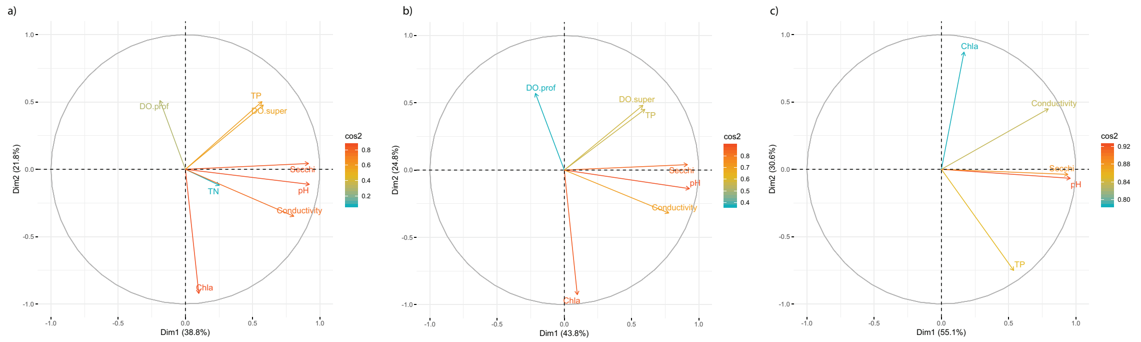


**Figure S4. Principal component analysis (PCA) results on water quality data.** (a) all collected historical water quality data; (b) collected historical water quality data excluding nitrates; (c) collected historical water quality data excluding nitrates, phosphates and chlorophyll-a.
